# Effect of 10 days of unilateral lower limb suspension on knee extensors neuromuscular function and spinal excitability

**DOI:** 10.1101/2024.03.06.583658

**Authors:** Loïc Lebesque, Marco V. Narici, Alain Martin, Giuseppe De Vito, Fabio Sarto, Gil Scaglioni

**Affiliations:** INSERM UMR1093-CAPS, Université Bourgogne Franche-Comté, UFR des Sciences du Sport, F-21000, Dijon; Department of Biomedical Sciences, University of Padova, Padova, 35131, Italy; CIR-MYO Myology Center, University of Padua, 35131, Italy

**Keywords:** Atrophy, Disuse, Force control, H-reflex, T-reflex, Unloading, Vastus medialis

## Abstract

The reduction in mechanical loading applied on the lower limb has numerous detrimental consequences on neuromuscular function. The current study aimed to investigate the changes in knee extensors’ strength and spinal excitability induced by unilateral lower limb suspension (ULLS), providing new insights into the neuromuscular adaptations to muscle hypoactivity. Ten young healthy males (19-28 years old) underwent 10 days of ULLS to simulate muscle disuse. Modulation by unloading of knee extensors’ function (muscle morphology and strength, activation capacity and contractile properties) and spinal reflexes were explored before and after the ULLS. The knee extensors’ anatomical cross-sectional area (−4%, *p* = 0.007), maximal strength (−27%, *p* < 0.001) and central activation ratio (−3%, *p* = 0.006) were reduced after 10 days of ULLS. Vastus medialis H-reflex amplitude was enhanced both at rest (+33%, *p* = 0.038) and during a low-intensity contraction set at 10% of maximal strength (+103%, *p* = 0.038). No changes in muscle contractility and nerve conduction velocity were observed after the ULLS. The present study suggests that neural impairments mainly contribute to the decrease in knee extensors’ strength induced by short-term ULLS. The decrease in muscle activation after a short period of ULLS was accompanied by an increase in spinal excitability. However, the latter adaptation did not counterbalance the reduction in activation capacity and thus in maximal strength resulting from ULLS. These adaptations to short-term ULLS should be considered when aiming at improving the neuromuscular function of people experiencing muscle hypoactivity.

**NEW & NOTEWORTHY:** This study provides new insights into the effects of muscle hypoactivity on neuromuscular function and spinal excitability in the major antigravity muscle group of the lower limb. Neural impairments primarily contribute to maximal strength loss after short-term unilateral lower limb suspension, while spinal excitability increased. These findings are crucial as they offer valuable understanding for developing effective interventions to improve health outcomes for individuals experiencing muscle inactivity.

## INTRODUCTION

Reductions in mechanical loading may occur as a consequence of surgery, injury and illness (1–3), but also as a result of reduced daily physical activity (4,5) or gravitational input, as during spaceflight (6). The direct outcome is a loss of muscle mass and function, associated with changes in motor control (7,8).

Several models of immobilization allow to explore the neuromuscular impairment in response to disuse (8). However, unilateral lower limb suspension (ULLS) enables to restrain atrophy to the explored musculoskeletal segment, easily mimicking the joint unloading that follows injuries or surgery (7,9) and, compared to casting, has the advantage of being less binding.

Extensive literature shows that the extent of muscle weakness is greater than the decrease in muscle mass since the first few days of ULLS, meaning that neuromuscular function is rapidly impaired by disuse (7,9–12). Among the antigravity muscles, the knee extensors appear to be the most affected by ULLS (13–15), showing a strength loss considerably larger than that of the plantar flexors (−1.0% · day^−1^ *vs.* 0,7% · day^−1^, for durations between 10 and 35 days of ULLS; 8,11). Therefore, although it is reasonable to assume that for unloading, as for overloading, changes in neural drive are the first to occur (8), their involvement in the weakening of antigravity muscles, associated with disuse, is still an unresolved issue. Although studies have shown that ULLS (21-30 days) does not interfere with the central motor drive of knee extensors muscles (9,11,15,16), others have found a decrease (17,18) or even an increase (range + 0.1%.day^−1^ increase to − 0.2%.day^−1^ decrease; median − 0.2%.day^−1^) of muscle voluntary activation (8). This divergence was also observed in surface electromyographic (EMG) data, recorded during maximal voluntary contractions, with studies showing a reduction of the vastii activity after ULLS (7-42 days) (19–23), and others finding no change for the vastii and RF (ULLS 7-35 days) (11,22,24). On the other hand, the literature agrees on the EMG data recorded during submaximal contractions. Interestingly, in this case, all studies showed that vastii EMG activity increased during submaximal contractions following ULLS (7,23–25), suggesting that additional motor units are necessary to produce the same absolute force output after unloading. In light of these considerations, it appears that changes at neural level have not yet been fully understood, and neither have the underlying mechanisms. Previous studies, on plantar flexors, have shown an unequivocal enhancement in the amplitude of the Hoffman (H)-reflex in response to unloading (7,8,12–14,26–30). The H-reflex is the EMG response evoked by electrically stimulating the Ia fibers of a peripheral nerve and reflects the efficacy of the synaptic transmission between Ia afferents and the motor neurons (31). Since it is generally considered as a comprehensive index of spinal excitability (e.g. 32,33), for greater readability we will use the term ‘spinal excitability’ to refer to the efficiency of the synaptic transmission between Ia afferents and motor neurons. The enhancement of H-reflex amplitude with unloading may be ascribed to an increased synaptic transmission efficiency and/or to a greater motoneuronal excitability (13,30,34), however, the improved synaptic transmission efficiency seems to be the predominant adaptation in all forms of disuse (13). This mechanism may interfere with the reduction in motoneuronal excitability that might occur together with an increased proportion of fast fibers in response to unloading (27). These adjustments should equally affect H and T-reflexes. The T-reflex is an electrophysiological variant of the stretch reflex caused by a mechanical tap applied on the tendon that, generating a quick stretch of the tendon itself, triggers muscle spindles discharge to Ia afferents notably. Both reflexes originate partly from the monosynaptic projections of the Ia afferents to alpha-motor neurones, but the T-reflex involves all the elements of the reflex pathway, including muscle spindles sensitivity and II afferents (35,36). The literature shows that the T-reflex may be further modulated by conflicting changes arising in peripheral components of the stretch reflex pathway. Actually, Anderson *et al.* (27) found, after 3 weeks of hind limb suspension in rat, a decline of the maximal soleus T-reflex but a rise in the maximal H-reflex (H_MAX_). Seynnes *et al.* (29) observed no change in soleus T-reflex, following 23 days of ULLS, but an increase in H-reflex and a decrease in tendon stiffness. The different modulations of the T-reflex response with unloading may be ascribed i) to a lower spindles’ sensitivity, due to their reduced solicitation during immobilization (37), and/or ii) to a decreased stiffness of the musculo-tendinous elements in series with the muscle spindles (27,29).

In view of these considerations and since the literature seems to show that few days of immobilization induce significant changes in neuromuscular function (8), this study sought to investigate the role of changes in spinal reflex excitability, muscle contractile characteristics and muscle morphology in the loss of knee extensors’ strength following 10 days of muscle unloading. Since, in plantar flexors, H-reflex is always modulated by ULLS and considering that knee extensors are among the antigravity muscles most affected by inactivity, we hypothesized that spinal excitability would be increased and muscle mass reduced by unloading. However, given the short period of immobilization, it could also be assumed that neural adaptations might be preponderant over muscular changes. Therefore, apart from improving knowledge on mechanisms responsible for the decline of the neuromuscular function due to unloading, the outcomes of the present investigation may be useful for refining the rehabilitation strategies of patients following periods of muscle disuse.

## MATERIALS & METHODS

### Ethical approval

The present study was approved by the Ethics Committee of the Department of Biomedical Sciences of the University of Padova (Italy) (reference number HEC-DSB/01-18) and conducted in accordance with the standards set by the latest revision of the Declaration of Helsinki. Participants were informed about all experimental procedures through an interview and an information sheet, and could ask questions to the selection committee. After signing the written consent form, participants were enrolled in the study with the possibility to withdraw at any stage of the experiment. This study was part of a larger investigation aiming to study the neuromuscular alterations with short-term disuse (18,38).

### Participants

Ten healthy and recreationally active young male volunteers (age: 22.9 ± 4.0 years old; body mass: 72.2 ± 7.2 kg; height: 177.6 ± 3.1 cm; body mass index: 22.9 ± 2.1 kg·m^−2^) took part to this study. Because unilateral lower limb suspension (ULLS) is associated with an increased risk of deep venous thrombosis (39), more commonly experienced in young females (40), only males were recruted in this study. Inclusion criteria were: age comprised between 18 and 35 years, body mass index comprised between 20 and 28 kg·m^−2^ and involvement in recreational physical activities 1-3 times per week. Exclusion criteria were: sedentary lifestyle, professional sport practice, smoking, history of deep venous thrombosis and acute or chronic musculoskeletal, metabolic and cardiovascular disorders. Using the Global Physical Activity Questionnaire (41), participants demonstrated a high level of physical activity (3268 ± 1468 MET·min·week^−1^) in the period leading up to the ULLS experiment. Volunteers were asked to maintain their habitual diet throughout the intervention period and refrain from intense exercise, coffee and alcohol intake during the 24 hours preceding the data collection at each time point.

### Protocol

#### Experimental design

Data were collected the day before (LS0) and after 10 days (LS10) of unilateral lower limb suspension (ULLS). For each subject, sessions lasted 2 hours and were planned at the same time of the day to avoid any circadian disturbance (42,43). At both LS0 and LS10, the experimental procedure was performed beginning with the evaluation of the maximal voluntary isometric contraction (MVC) of knee extensors and the maximal voluntary activation level (VAL). Then, vastus medialis (VM) spinal excitability were assessed through electrical stimulations and patella tendon striking at rest and during a weak contraction of knee extensors (10% MVC assessed for each session). In addition, quadriceps and VM morphology was assessed through ultrasonography, on the same day of the ULLS beginning (baseline data collection, BDC) and the day before the end of the ULLS (LS9).

#### Maximal force production capacity

To start the maximal force production evaluation, subjects performed a knee extensors warm-up composed of 10 submaximal contractions, lasting 4 s each. Then, three knee extensors MVCs (lasting 4 s, with 1-min rest in-between) were performed with maximal electrical doublet (100 Hz) delivered over the peak force.

#### Spinal excitability

Since it is more challenging to evoke rest H-reflex in the quadriceps muscles compared to the soleus muscle, several studies suggest eliciting it during a weak muscle contraction (44–47). The reflex response is indeed enhanced by the muscle contraction because motoneurons are brought closer to their firing threshold by the voluntary drive (48). In addition, muscle activation allows to keep constant the “background” level of α-motoneuron depolarisation and limits post-synaptic modulations (45,49). Therefore, the assessment of spinal excitability during muscle contraction has the advantage of increasing amplitude and reducing variability in both latency and amplitude of the H-reflex (48,50,51). Consequently, VM H-reflex and M-wave recruitment curves were built by applying electrical stimulations first during weak knee extensors contractions (10% MVC) and then at rest. To build the recruitment curves, stimulations started at 5 mA and were progressively increased with a 5-mA increment until the maximal M-wave (M_MAX_) was reached. To ensure that the M-wave lied in the plateau of its maximal value throughout the experiment, the maximal stimulation intensity was increased by 20%. Subsequently, additional stimulations were applied, in 2 mA increments, around the intensity that allowed to record the highest H-reflex, in order to accurately estimate the intensity needed to elicit the maximal H reflex (H_MAX_). Four stimuli were delivered at each intensity interspaced by 10 s intervals. Then, 6 out of 10 subjects underwent complementary evaluation of spinal excitability at rest through the assessment of tendinous reflex (i.e., T-reflex, for the method see below mechanical stimulation).

#### Unilateral lower limb suspension

The ULLS model was employed for 10 days. The non-dominant leg (left leg for all participants) of the participants was fitted with a shoe having an elevated sole (50 mm), while the dominant leg (right leg for all participants) was suspended and kept at a slightly flexed position (approximately 15-20° of knee flexion) using straps. Volunteers were asked to walk on crutches during the whole ULLS period and to refrain from loading the suspended leg in any way. A familiarization session was set up for the participants to practice daily tasks with the limb suspended (9). Participants were recommended to wear elastic compression socks on the suspended leg during ULLS period and to perform passive, range of motion, non-weight-bearing exercises of the ankle, as precautionary measures to prevent deep venous thrombosis (39). Moreover, we performed an ultrasound-Doppler examination after 5 days of ULLS. Participants’ compliance was evaluated through daily calls and messages and by comparing calves’ temperature and circumference after 5 and 10 days of ULLS, as previously suggested (9).

### Data acquisition

#### Ultrasonography

The morphology of the quadriceps was assessed by ultrasonography, which also allowed us to define the morphology of the VM, for which electrophysiological data were also recorded. To evaluate morphology of quadriceps and VM muscles, subjects were lying on their backs on an auscultation table. Muscles anatomical cross-sectional area (ACSA) was obtained using extended-field-of-view ultrasonography imaging (Mylab, Esaote, Genoa, Italy). A 47 mm, 7.5 MHz linear array transducer was used to collect images at different muscle length percentages. Three regions of interest were detected at 30%, 50% and 70% of femur length (measured as the distance between the great trochanter and the mid-patellar point), where 0% represents the mid-patellar point (distal part) and 100% the greater trochanter (proximal part). An adjustable guide was used in each acquisition to keep the same transverse path (52). After applying a generous amount of transmission gel on the participant’s skin to improve the acoustic contact, the transducer was moved slowly in the transverse plane from the medial border of the VM to the lateral border of the vastus lateralis. During the transducer displacement, we kept the pressure on the skin as constant as possible. Two scans were obtained for each site.

#### Force recordings

Voluntary and electrically-evoked force of knee extensors was recorded using a custom-made knee dynamometer fitted with a strain gauge. The hip and knee angles were set at 90° (180° full extension). Subject’s dominant leg was securely strapped to the custom-made knee dynamometer, equipped with a load cell, at the ankle level. Subject was asked to cross the arms on the chest and keep the head in a neutral position during the experiment procedure. The force exerted by the subject was displayed in real-time on a computer screen placed at 1 m in front of him. Because spinal excitability was further assessed during a weak quadriceps contraction, a line representing 10% of the session MVC was added on the computer screen once the MVC was determined. Force data were were recorded at a sampling frequency of 1 kHz with the Spike2 software (version 8.01, Cambridge Electronics Design Ltd, Cambridge, UK; RRID: SCR_000903) and stored for offline analysis with the LabChart software (version 8, AD Instruments®, Australia; RRID: SCR_017551).

#### Electromyographic recordings

Electromyographic activity (EMG) from the dominant-leg VM was recorded during the 10% MVC contractions with bipolar active-type Ag-AgCl electrodes electrodes (44 x 22 mm, Ambu®). To minimize skin impedance (< 5 kΩ), the skin was shaved, rubbed with an abrasive paste and cleaned with alcohol, before electrode positioning. Electrodes were positioned 10 cm above the patella upper part and 4 cm medially with an inter-electrode center-to-center distance of 20 mm. The EMG signals were amplified (gain = 1,000) and bandpass filtered at 5 Hz to 5 kHz using a CED 1902 amplifier (Cambridge Electronics Design Ltd, Cambridge, UK) and digitized with a CED Micro 1401 data acquisition unit (Cambridge Electronics Design Ltd, Cambridge, UK). EMG data were recorded at a sampling frequency of 40 kHz with the Spike2 software (version 8.01, Cambridge Electronics Design Ltd, Cambridge, UK; RRID: SCR_000903) and stored for offline analysis with the LabChart software (version 8, AD Instruments®, Australia; RRID: SCR_017551).

#### Nerve electrical stimulations

Nerve electrical stimulations were delivered using a high-voltage (400 V) constant-current stimulator (DS7AH model, Digitimer, Hertfordshire, UK). For maximal voluntary activation assessment, electrical doublets (100 Hz, monophasic rectangular pulses, 1 ms width) were delivered through two stimulation electrodes (5 x 10 cm, Compex SA, Ecublens, Switzerland) placed in the proximal and distal region of the knee extensors. Maximal electrical stimulation intensity was determined as the intensity eliciting the maximal twitch amplitude. For spinal excitability assessment, we delivered percutaneous electrical stimulations (monophasic rectangular pulses, 1 ms width) in the femoral trigone over the femoral nerve to evoke electrophysiological and mechanical responses of the VM muscle. The self-adhesive anode (5 x 10 cm, Compex SA, Ecublens, Switzerland) was placed in the gluteal fossa. The optimum stimulation site was located with a hand-held cathode stylus in order to obtain the greatest H-reflex amplitude of the VM for the lowest stimulation intensity. Then, the self-adhesive cathode electrode (7 mm diameter, Contrôle Graphique Medical, Brie-Compte-Robert, France) was positioned and firmly fixed with a velcro elastic band.

#### Mecanichal stimulations

During T-reflex procedure, subjects were positioned as described previously in *Force recordings* section. Twenty T-reflexes were elicited for each subject by tapping the anterior face of the patellar tendon of the quadriceps with a manual reflex hammer (neurological hammer, Babinsky model) at various intensities (53). The T-reflex on the VM EMG signal and its associated mechanical twitch were recorded for each strike.

### Data analysis

#### Ultrasonography

Of the two scans obtained at each site, we analyzed the image with the best quality. Quadriceps and VM ACSAs were measured by tracing the muscle contours with the image analysis software “ImageJ” (https://imagej.net; RRID: SCR_003070). Two measurements per parameter were performed on each image, and their average was considered. Quadriceps ACSA (qACSA) was computed as the average value of the ACSAs recorded at 30, 50 and 70% of the femur length for each subject at both LS0 and LS10. VM ACSA was calculated as the average value of the ACSAs recorded at 30 and 50% of the femur length, due to technical issues in clearly detecting the border between the VM and vastus intermedius at 70% of the femur length.

#### Maximal force production

The MVC was determined as the highest force value reached during the 3 MVCs. The target of 10% MVC was calculated from the MVC of each session (i.e., LS0 or LS10). The normalized force was calculated as the MVC divided by the quadriceps ACSA (MVC/qACSA).

#### Activation capacity

The voluntary activation associated with the maximal force production and the weak contraction (10% MVC) was quantified with the central activation ratio (CAR_MVC_ and CAR_10%_ respectively) as follows (54):

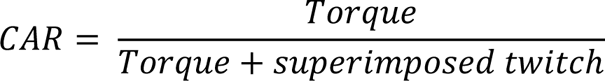

where torque was either the MVC (CAR_MVC_) or the 10% MVC (CAR_10%_). A higher CAR value indicates an increase in muscle activation. It should be noted that, while the CAR_MVC_ was calculated using maximal doublet stimulations (100 Hz) delivered during brief MVC, the CAR_10%_ was derived from the M-wave recruitment curve recorded during weak contractions at 10% MVC using maximal single stimulations (see above *Spinal excitability* section). The ratio between the CAR_10%_ and the force produced during the contraction at 10% MVC was calculated to determine the motor unit activity required to achieve this specific target force. The root mean square (RMS) of VM EMG activity was computed offline over a 500 ms period preceding H_MAX_ and M_MAX_ stimulations during the weak contractions at 10% MVC, and subsequently normalized to the M_MAX_ amplitude (RMS/M_MAX_).

#### Spinal excitability parameters

One subject has been exempted from the spinal excitability assessment because he did not tolerate electrical stimulations over the femoral nerve. Peak-to-peak amplitude of both H-reflex and M-wave was measured and averaged for each stimulation intensity. The highest amplitude of M-wave and H-reflex were labelled H_MAX_ and M_MAX_ respectively. The H_MAX_/M_MAX_ ratio was then calculated. In one subject it was not possible to evoke the H-reflex at rest and was excluded from H_MAX_-related parameters analysis at rest. The amplitude of the M-wave accompanying H_MAX_ (M_atHMAX_) was also measured and normalized by M_MAX_ to verify that stimulation conditions were similar between sessions. The RMS of VM EMG signal was measured over the 500 ms preceding each H_MAX_ (RMS_H_) and M_MAX_ (RMS_M_) electrical stimulation during the contraction. We calculated the H_MAX_/RMS_H_ ratio to assess the influence of muscle activity background on spinal excitability (55,56). We also measured H_MAX_ and M_MAX_ latencies at rest (H_LATENCY_ and M_LATENCY_) as the delay between the onset of stimulation artefact and the upshot of electrophysiological response. While M_LATENCY_ represents the time required for conduction down the motor axons to the terminal branches and propagation across the neuromuscular junction, H_LATENCY_ reflects the time required for signal propagation through the reflex arc, including the Ia afferent synaptic delay at the spinal level. Moreover, the difference between H_LATENCY_ and M_LATENCY_ (H-M_LATENCY_), representing an index of nerve conduction velocity through the spinal reflex loop, was computed (13). Then, these latencies were normalized by the subject’s femur length (i.e., the distance between the greater trochanter and the lateral condyle). For one subject, the onset of the VM H_MAX_ was merged with the associated M-wave, thus this subject was excluded from H_MAX_ latency analysis. The amplitude of the mechanical twitch associated with both M_MAX_ (M_TW_) and H_MAX_ at rest was measured. Since the M_TW_ represents the mechanical response of the muscle when the maximum amount of the motor units is recruited (57), its characteristics were analyzed. For each M_TW_, we determined the contraction time (i.e., the time between twitch onset and maximal twitch value; CT), the rate of force development (i.e., the maximal value of the first derivative of the force signal using a 1 ms sliding window; RFD) and the half relaxation time (i.e., corresponding to the time needed to reach a 50% decrease of twitch force; HRT) before computing the averages of the four measurements. To assess the electromechanical efficiency, the M_TW_/M_MAX_ was calculated. We analyzed the relationship between the amplitude of the T-reflex and the corresponding twitch since we did not have the complete T-reflex recruitment curve and therefore were not certain of obtaining the maximal T-wave response. The relationship between VM T-reflex amplitudes, expressed as a percentage of M_MAX_, and the associated twitch amplitudes were scatter-plotted individually. The slope and the intercept of the linear regression of these relationships were determined for each subject and averaged at LS0 and LS10.

#### Statistical analysis

The normality of each dataset was assessed with the Shapiro-Wilk test. For all variables that passed normality test, Student’s paired t-test was applied in order to investigate whether difference between LS0 and LS10 was present within the considered parameters. When normality test was not passed, non-parametric Wilcoxon’s signed-rank test for paired comparison was used. Only for the RMS/M_MAX_ variable, a two-way repeated measures ANOVA with *time* (LS0 *vs*. LS10) and *stimulation* (H_MAX_ *vs.* M_MAX_) factors was performed. If a significant effect was found, a Tukey’s post-hoc analysis was conducted. Correlations between MVC and qACSA as well as between MVC and CAR_MVC_ were analyzed using repeated measures correlations (rmcorr R package, 51). Effect sizes (ES) were calculated with Cohen’s d and r respectively for parametric and non-parametric comparison (59). Small, moderate and large effects are considered for Cohen’s d ≥ 0.2, ≥ 0.5, ≥ 0.8, and Cohen’s r ≥ 0.1, ≥ 0.3, ≥ 0.5 respectively. Results are reported as mean ± standard deviation. Significance level was set at *p* < 0.05 and the confidence interval at 95% (CI_95%_). Student’s and Wilcoxon’s tests were performed using Statistica software (Statsoft, version 12, Tulsa, OK, USA; RRID: SCR_014213) while Cohen’s effect sizes were calculated with G*Power software (version 3.1.9.2, Universität Düsseldorf, Germany; RRID: SCR_013726). Repeated measures correlations were performed with R (version 4.1.3, R Core Team, 2017; RRID: SCR_001905). Graphs were made with GraphPad Prism software (version 8.4.0, GraphPad software Inc. San Diego, CA, USA; RRID: SCR_002798).

## RESULTS

### Muscle morphology, strength and activation capacity

The ACSA of the unloaded VM (t_(9)_ = 4.042, CI_95%_ = [0.48 : 1.69], *p* = 0.003, ES = 0.426) and quadriceps (t_(9)_ = 3.444, CI_95%_ = [0.87: 4.19], *p* = 0.007, ES = 0.397) decreased (−7.8 and -4.4 %, respectively) after 10 days of ULLS (Table 1). The ACSA losses were accompanied by a 27.1 % reduction in knee extensors MVC (t_(9)_ = 9.464, CI_95%_ = [17.01 : 27.70], *p* < 0.0001, ES = 2.196) (Table 1). The normalized force (MVC/qACSA) was 23.7 % lower after ULLS (t_(9)_ = 8.792, CI_95%_ = [0.25 : 0.43], *p* = 0.0001, ES = 2.016) (Table 1).

**Table 1.** Muscle morphology, strength and activation capacity.

| | BDC ($n = 10$) | LS9 ($n = 10$) | *p*-value |
| --- | --- | --- | --- |
| VM ACSA ($\text{cm}^2$) | $16.34 \pm 2.78$ | $15.26 \pm 2.26$ | 0.003 |
| qACSA ($\text{cm}^2$) | $57.79 \pm 6.54$ | $55.26 \pm 6.20$ | 0.007 |
| | LS0 ($n = 10$) | LS10 ($n = 10$) | *p*-value |
| MVC (kg) | $82.54 \pm 11.13$ | $60.19 \pm 9.13$ | $< 0.0001$ |
| 10% MVC (kg) | $8.25 \pm 1.11$ | $6.02 \pm 0.91$ | $< 0.0001$ |
| MVC/qACSA ($\text{kg} \cdot \text{cm}^{-2}$) | $1.43 \pm 0.18$ | $1.09 \pm 0.16$ | 0.0001 |
| CAR<sub>MVC</sub> | $0.980 \pm 0.013$ | $0.950 \pm 0.033$ | 0.006 |
| CAR<sub>MVC</sub>/MVC ($\text{kg}^{-1}$) | $0.012 \pm 0.002$ | $0.016 \pm 0.002$ | $< 0.0001$ |
| CAR<sub>10%</sub> | $0.428 \pm 0.065$ | $0.364 \pm 0.063$ | 0.003 |
| CAR<sub>10%</sub>/10% MVC ($\text{kg}^{-1}$) | $0.052 \pm 0.009$ | $0.061 \pm 0.011$ | 0.032 |
| RMS/$M_{MAX}$ at 10% MVC<br>($\times 10^{-3}$) | $9.39 \pm 4.05$ | $6.18 \pm 3.54$ | 0.020 |

All variables were measured before (LS0) and after (LS10) 10 days of unilateral lower limb suspension, except ACSA measured the day of ULLS beginning (BDC) and the day before the ULLS end (LS9). MVC = knee extensors maximal voluntary contraction; VM = vastus medialis muscle; ACSA = muscle anatomical cross-sectional area; qACSA = quadriceps ACSA; CAR<sub>MVC</sub> and
CAR<sub>10%</sub> = central activation ratio calculated during a MVC and during a contraction at 10% MVC; RMS = root mean square; M<sub>MAX</sub> = maximal M-wave. Values are reported as mean ± standard deviation.

Activation capacity as measured by CAR_MVC_ decreased by 3.0 % after ULLS (t_(9)_ = 3.557, CI_95%_ = [0.01 : 0.05], *p* = 0.006, ES = 1.175) (Figure 1) while the CAR_10%_ decreased by 15.0 % (t_(9)_ = 4.011, CI_95%_ = [2.79 : 10.03], *p* = 0.003, ES = 0.997). The ratio between the CAR_MVC_ and the MVC increases by 33.4 % following ULLS (t_(9)_ = -6.839, CI_95%_ = [-0.01 : -0.00], *p* < 0.0001, ES = 1.880), while the ratio between the CAR_10%_ and the force produced at 10% of MVC increased by 16.5 % (t_(9)_ = -2.540, CI_95%_ = [-0.02 : -0.00], *p* = 0.032, ES = 0.869) (Table 1). Furthermore, the normalized EMG activity (RMS/M_MAX_) of the VM during the 10% MVC plateau declined by 34.2% after ULLS (only *time* effect; F(_1,8_) = 8.482, *p* = 0.020, *η*_p_^2^ = 0.515 ; Tukey post-hoc : *p* = 0.020, ES = 0.839) (Table 1). Based on repeated measures correlation analysis, it appears that MVC is positively correlated to qACSA (r_rm_ (9) = 0.773, CI_95%_ = [0.22 : 0.95], *p* = 0.005) (Figure 2A) and CAR_MVC_ (r_rm_ (9) = 0.624, CI_95%_ = [-0.07 : 0.91], *p* = 0.040) (Figure 2B). In other words, the higher the qACSA or CAR_MVC_, the greater the production of maximal force. However, MVC change was not correlated to changes in CAR_MVC_ (r = -0.382, r² = 0.146, *p* = 0.276) and qACSA (r = 0.189, r² = 0.036, *p* = 0.600) (see Supplemental Figure S1).

**Figure 1.**
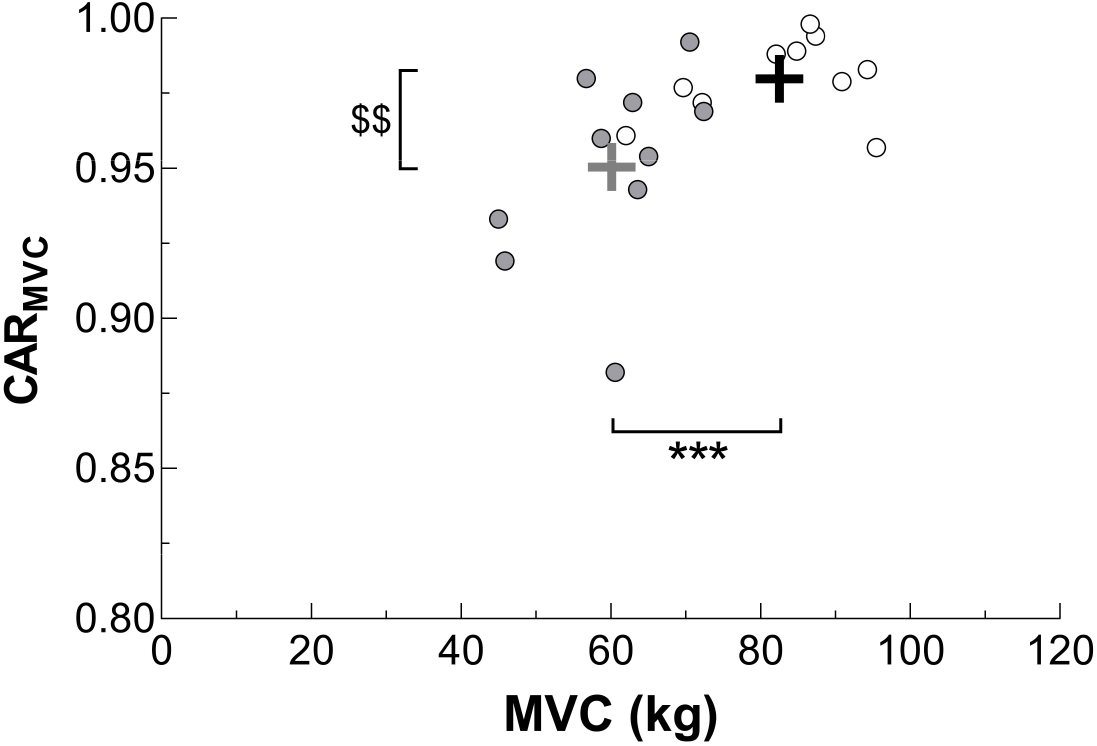
Maximal force and corresponding central activation. The maximal voluntary contraction (MVC) and corresponding central activation ratio (CAR_MVC_) were individually scatter plotted before (white circles) and after (grey circles) 10 days of unilateral lower limb suspension (*n* = 10). Black (before limb suspension) and grey (after limb suspension) crosses represent the mean of both scatter plots. \*\*\**p* < 0.001 and ^$$^*p* < 0.01 express significant differences in MVC and CAR_MVC_, respectively, between before and after lower limb suspension.

**Figure 2.**
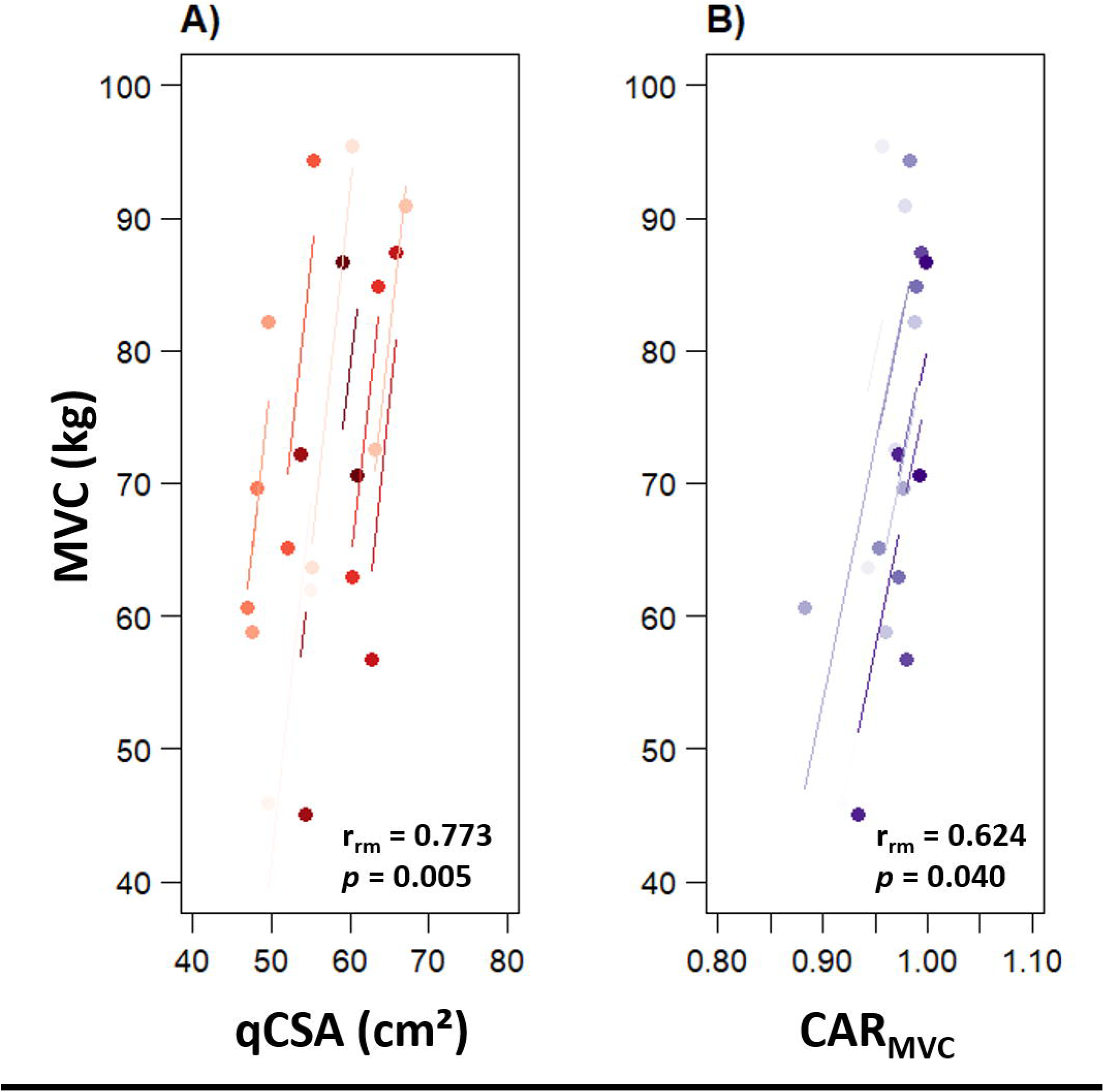
Individual repeated measures correlations (rmcorr) between muscle function parameters and maximal force production (*n* = 10). Rmcorr was plotted between qACSA and MVC (A), and between CAR and MVC (B). Observations from the same participant are given the same colour, the lines correspond to the rmcorr fit for each participant. MVC = maximal voluntary contraction; qACSA = quadriceps cross-sectional area; CAR = central activation ratio.

### Muscle Contractile Properties

Mechanical parameters associated with H_MAX_ and M_MAX_ stimulations at rest were also compared between LS0 and LS10. The ULLS did not modify the twitch amplitude associated with M_MAX_ (M_TW_; t_(8)_ = 1.593, CI_95%_ = [-0.69 : 3.75], *p* = 0.150, ES = 0.599) (Figure 3). Specifically, the RFD of M_TW_ was not altered after 10 days of ULLS (t_(8)_ = 1.685, CI_95%_ = [-0.01 : 0.09], *p* = 0.130, ES = 0.678), nor were the CT (t_(8)_ = -1.159, CI_95%_ = [-14.61 : 4.84], *p* = 0.280, ES = 0.420) and the HRT (t_(8)_ = 0.165, CI_95%_ = [-15.05 : 17.36], *p* = 0.873, ES = 0.070). Also, the M_TW_/M_MAX_ ratio was not changed by the ULLS protocol (Z = 0.770, *p* = 0.441, ES = 0.257). Muscle contractile properties obtained at rest are provided in Table 2. Regarding the twitch associated with H_MAX_, its amplitude was not different between LS0 and LS10 (t(_7_) = 1.015, CI_95%_ = [-0.84 : 2.11], *p* = 0.344, ES = 0.253) (Figure 3).

**Figure 3.**
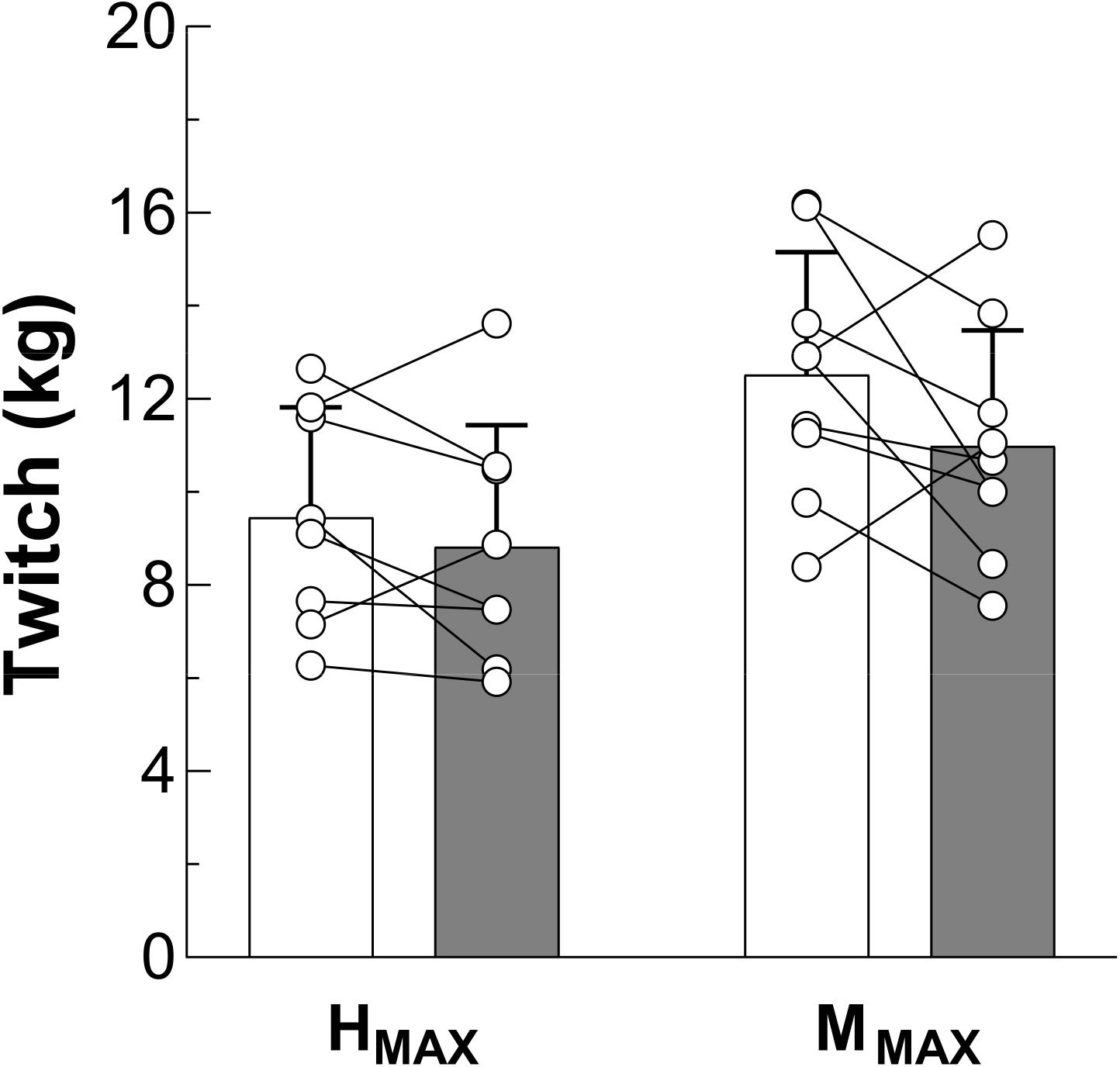
Twitch amplitude associated with H_MAX_ and M_MAX_. The twitch amplitude associated with H_MAX_ (*n* = 8) and M_MAX_ (*n* = 9) was measured at rest before (white columns) and after (grey columns) 10 days of unilateral lower limb suspension. Circles and lines represent individual data evolution. H_MAX_ = maximal H-reflex; M_MAX_ = maximal M-wave.

**Table 2.** Mechanical parameters associated with M_MAX_ before and after ULLS.

|  | M<sub>MAX</sub> stimulations |  | *p*-value |
| --- | --- | --- | --- |
|  | LS0 (*n* = 9) | LS10 (*n* = 9) |  |
| M<sub>TW</sub>/M<sub>MAX</sub> (kg/mV) | 1.38 ± 1.07 | 1.33 ± 1.25 | 0.441 |
| CT (ms) | 82.44 ± 8.59 | 87.33 ± 14.03 | 0.280 |
| RFD (kg/ms) | 0.247 ± 0.060 | 0.210 ± 0.048 | 0.130 |
| HRT (ms) | 83.60 ± 15.15 | 82.44 ± 17.80 | 0.873 |

M<sub>MAX</sub> = maximal M-wave; M<sub>TW</sub> = twitch associated with M<sub>MAX</sub>; CT = twitch contraction time; RFD = twitch rate of force development; HRT = twitch half relaxation time; ULLS = unilateral lower limb suspension; LS0 and LS10 = before and after 10 days of ULLS. Values are reported as mean ± standard deviation.

### H-Reflex responses and M-wave

While the H_MAX_ at rest was increased by 59.0% after 10 days of ULLS (Z = 2.100, *p* = 0.036, ES = 0.743), H_MAX_ obtained during muscle contractions at 10% MVC was not modified (Z = 0.770, *p* = 0.441, ES = 0.257) (Table 3). It should be noted that the amplitude of the M-wave associated with H_MAX_ (M_atHMAX_) was not different between LS0 and LS10 either at rest (Z = 0.700, *p* = 0.484, ES = 0.247) or during contraction (Z = 1.362, *p* = 0.173, ES = 0.454) (Table 3). Therefore, the stimulation conditions at H_MAX_ may be considered stable, allowing the comparison of spinal excitability features between the two time points. M_MAX_ did not differ between LS0 and LS10 either at rest (t_(8)_ = -0.258, CI_95%_ = [-1.83 : 1.46], *p* = 0.803, ES = 0.025) or during 10% MVC contraction (t_(8)_ = 1.345, CI_95%_ = [-0.42 : 1.59], *p* = 0.215, ES = 0.074) (Table 3). The ULLS increased the H_MAX_/M_MAX_ at rest by 33.1 % (LS0 : 0.20 ± 0.15 *vs*. LS10 : 0.27 ± 0.15 ; t(_7_) = -2.557, CI_95%_ = [-0.13 : -0.01], *p* = 0.038, ES = 0.441) but had no impact on the ratio during contraction at 10% MVC (LS0 : 0.34 ± 0.16 *vs.* LS10: 0.36 ± 0.17; t_(8)_ = -1.947, CI_95%_ = [-0.04 : 0.00], *p* = 0.087, ES = 0.121) (Figure 4A). This result is also depicted in Figure 4B where experimental traces of M_MAX_ and H_MAX_ obtained at rest, before and after ULLS, from one representative subject, are presented. In order to overcome differences between sessions in EMG activity during contractions at 10% MVC, we also normalized H_MAX_ by muscle activity background preceding H_MAX_ stimulations (RMS_H_), and we observed an increase of this ratio after the ULLS period (+102.9 % ; t_(8)_ = -2.480, CI_95%_ = [-74.00 : -2.70], *p* = 0.038, ES = 1.097) (Table 3 and Figure 4C).

**Figure 4.**
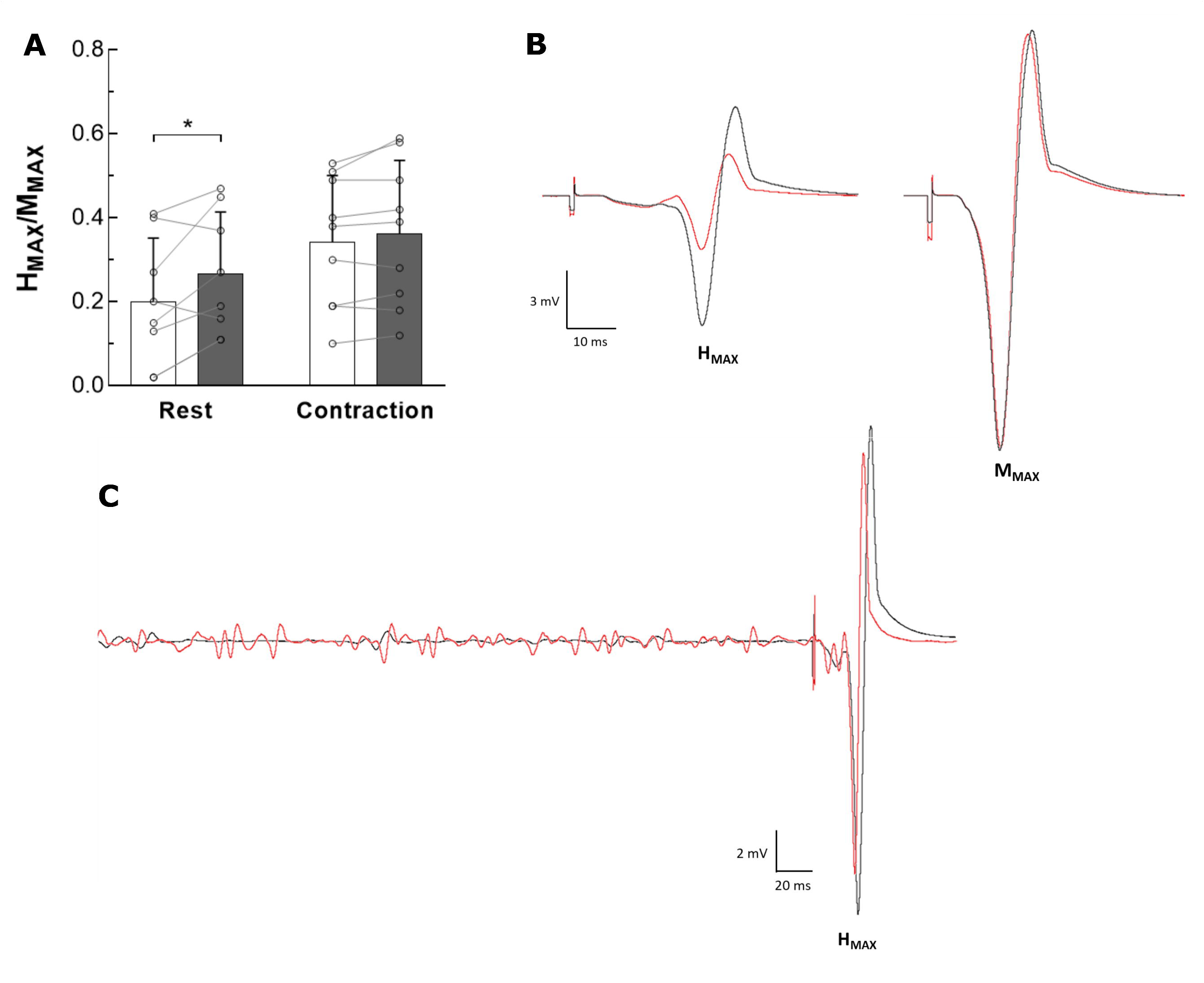
Changes in spinal excitability after 10 days of ULLS. A. H_MAX_/M_MAX_ ratio was calculated before (white columns) and after (grey columns) 10 days of ULLS, either at rest (*n* = 8) and during 10% MVC contractions (*n* = 9). Circles and lines represent individual data evolution. B. H_MAX_ and M_MAX_ traces at rest, from a typical subject, before (red lines) and after (black lines) ULLS. C. Representative data of VM EMG activity during the 10% MVC contraction with associated H_MAX_ before (red line) and after (black line) ULLS from a typical subject. VM EMG = electromyographic activity of vastus medialis; H_MAX_ = maximal H-reflex; M_MAX_ = maximal M-wave. ULLS = unilateral lower limb suspension. *\**p < 0.05 express significant difference.

**Table 3.** Spinal excitability and nerve conduction parameters before and after 10 days of ULLS.

|  | Rest |  |  | Contraction |  |  |
| --- | --- | --- | --- | --- | --- | --- |
|  | LS0 | LS10 | *p*-value | LS0 | LS10 | *p*-value |
| M<sub>atHMAX</sub> (% M<sub>MAX</sub>) | 8.39 ± 7.35 | 6.64 ± 8.65 | 0.484 | 9.02 ± 9.03 | 5.17 ± 4.63 | 0.173 |
| H<sub>MAX</sub> (mV) | 2.22 ± 1.94 | 3.53 ± 3.25 | 0.036 | 4.29 ± 3.21 | 4.37 ± 3.44 | 0.441 |
| M<sub>MAX</sub> (mV) | 13.12 ± 7.04 | 13.30 ± 7.41 | 0.803 | 13.48 ± 7.97 | 12.89 ± 7.91 | 0.215 |
| H<sub>MAX</sub>/RMS<sub>H</sub> | - | - | - | 37.27 ± 12.95 | 75.62 ± 47.69 | 0.038 |
| M<sub>LATENCY</sub> (ms/m) | 15.22 ± 1.60 | 15.74 ± 1.81 | 0.613 | - | - | - |
| H<sub>LATENCY</sub> (ms/m) | 48.04 ± 5.55 | 48.72 ± 5.55 | 0.396 | - | - | - |
| H-M<sub>LATENCY</sub> (ms/m) | 34.07 ± 5.34 | 33.84 ± 4.02 | 0.782 | - | - | - |
| T-twitch slope | 0.253 ± 0.096 | 0.331 ± 0.275 | 0.422 | - | - | - |
| T-twitch intercept | 0.876 ± 0.585 | 1.550 ± 0.807 | 0.221 | - | - | - |

Data were obtained at rest and/or during 10% MVC contractions. For each parameter, the number of subjects was 9, except for M<sub>atHMAX</sub> at rest (*n* = 8), H<sub>LATENCY</sub> and H-M<sub>LATENCY</sub> (*n* = 7), T-
twitch slope and intercept ( $n = 6$ ). LS0 = before leg suspension; LS10 = after 10 days of leg suspension; $H_{MAX}$ = maximal H-reflex; $M_{MAX}$ = maximal M-wave ; $M_{atHMAX}$ = M-wave associated with maximal H-reflex; $H_{LATENCY}$ and $M_{LATENCY}$ = $H_{MAX}$ and $M_{MAX}$ latency respectively; $H-M_{LATENCY}$ = difference between $H_{LATENCY}$ and $M_{LATENCY}$ ; T = T-reflex; twitch = T-reflex associated twitch; MVC = maximal voluntary contraction; ULLS = unilateral lower limb suspension. Values are reported as mean ± standard deviation. Please note that some parameters are obtained only in one condition (rest or contraction). $*p < 0.05$ and $**p < 0.01$ express significant difference from LS0 for the considered condition. Values are reported as mean ± standard deviation.

Moreover, the delay between electrical stimulation and electrophysiological responses was determined for the M_MAX_ and H_MAX_ waves. The latency of M_MAX_ (M_LATENCY_) and H_MAX_ (H_LATENCY_) was not affected by ULLS (M_LATENCY_ : t_(8)_ = -0.527, CI_95%_ = [-1.75 : 1.10], *p* = 0,613, ES = 0.192; H_LATENCY_ : t(_6_) = -0.913, CI_95%_ = [-2.53 : 1.15], *p* = 0,396, ES = 0.134) (Table 3). The difference between H_LATENCY_ and M_LATENCY_ (H-M_LATENCY_) was not changed after 10 days of ULLS (t(_6_) = 0.289, CI_95%_ = [-1.73 : 2.20], *p* = 0.782, ES = 0.049).

In addition, the ULLS did not modulate the relationship between the VM T-reflex and the associated twitch. Indeed, the unloading period did not alter the slope (t(_5_) = -0.874, CI_95%_ = [-0.31 : 0.15], p = 0.422, ES = 0.381) and the intercept (t(_5_) = -1.397, CI_95%_ = [-1.91 : 0.57], *p* = 0.221, ES = 0.946) of this linear regression (Table 3).

## DISCUSSION

The present study aimed to investigate neural, contractile and morphological changes associated with muscle weakness following 10 days of ULLS of the main antigravity muscle group of the lower limbs. We showed that knee extensors’ disuse led to a reduction in MVC mainly explained by an impairment in central motor drive. For the first time, spinal excitability was investigated in the quadriceps muscle group to deepen knowledge of neural adaptation to unloading in antigravity muscles. Our main results show that spinal excitability was enhanced at rest and during muscle contraction following 10 days of ULLS. These findings seem particularly relevant as they provide new insights into the neurophysiological adjustments that occur in a major antigravity muscle in response to human muscle disuse.

### Muscle morphology and function

The present investigation shows that 10 days of ULLS reduced the maximal force developed by the knee extensors (−27 %, -2.7 %·day^−1^). This alteration in MVC was associated with a reduction in qACSA of 0.44 %·day^−1^ (−4.4 % in 10 days). In overall our results are consistent with previous investigations that reported an impairment in force production capacity (11,14–16,23,25,60) as well as a decrease in knee extensors size after ULLS (8,11,16,23)of the knee extensors . As reported in the literature, muscle atrophy could be explained by decreases in protein synthesis rate after lower limb muscle disuse (61,62) affecting the muscle protein synthesis/breakdown balance (63) and thus, the myofibrillar content (64). The present study corroborated, in agreement with earlier investigations (8,16,65–67), that strength loss, after unloading, was greater than the rate of muscle size loss, therefore resulting in an alteration of normalized force (−24 %). It is thought that muscular and tendinous alterations are involved, such as a reduction in myosin content (16,68) and tendon stiffness (11), and an impairment of the excitation-contraction coupling (67). Neural alterations, including a reduction in voluntary activation and coordination of antagonistic muscles (69), could also be potential explanatory factors contributing to the impairment of normalized force during muscle disuse. In the current study, we observed an impairment of both maximal and submaximal central motor drive, as revealed by the decrease in CAR_MVC_ (−3 %), CAR_10%_ (−15 %) and RMS/M_MAX_ (−34 %). Particularly, a decrease in CAR_MVC_ was associated with a reduction in maximal force production capacity. In addition, our data showed a rise in CAR_MVC_/MVC and CAR_10%_/10% MVC ratios after the ULLS protocol, suggesting an impairment in muscle electromechanical efficiency (i.e. the dissociation between electrical and mechanical events of muscle function; 68) after knee extensors unloading. Though our findings are consistent with some previous investigations on knee extensors and plantar flexors after 21-30 days of limb suspension (23,29,60,71), some others did not observed any change in maximal muscle activation, for the same muscle groups, after 10-30 days of limb unloading (11,13,16,25,30,72). Despite these conflicting results, strength loss after unloading seems to be more related to alterations in neural than muscular factors (13,73). As a matter of fact, among multiple predictive variables, including muscle ACSA, twitch RFD and HRT or the amplitude of the H-reflex and M-wave for instance, it appears that central activation is the most influential variable in mediating strength loss after muscle disuse (13).

Regarding mechanical twitch evoked by H_MAX_ stimulation at rest, no change was reported following 10 days of ULLS in the current study. The present observation contrasts with the result of Seynnes *et al.* (29) which reported a 40 and 54% increase in H_MAX_ twitch amplitude for the plantar flexors after 14 and 23 days of ULLS respectively. This discrepancy might depend on the muscle group investigated and the different duration of exposure to limb unloading. On the other hand, in agreement with data from Campbell *et al.* (16) following 21 days of ULLS, the current analysis, did not find any change in M_TW_. However, it should be noted that there is no consensus in the literature about the evolution of M_TW_ following muscle disuse, since studies have shown that ULLS decreased (15,60,65) or even increased (30) the maximal twitch amplitude on lower limb muscles. In the current study, the M_TW_ parameters (CT, RFD and HRT) were unaltered after the knee extensors’ unloading. These outcomes are in accordance with the literature focusing on the muscle contractility of knee extensors following 21-30 days of ULLS (16,30,60). Specifically, lower limb suspension did not result in the alteration of M_TW_ CT (16), RFD (16,30,60) and HRT (16,30,60). It should be noted that we evoked a twitch to assess muscular factors of knee extensors. The sensitivity of this index could not be sufficient to detect changes in muscle contractility (74). Despite this, the present findings underline the discrepancy between morphological (i.e. reduction in qACSA, maximal strength) and functional (i.e. no change in quadriceps contractility) data already observed in the literature (8) and suggest that short-term ULLS-induced strength loss is primarily attributed to neural factors. Therefore, the present model of disuse does not seem to impair knee extensors’ contractility and relaxation processes, although it is effective in inducing strength loss through impairments in muscle activation. Due to the relatively small number of data obtained for muscle contractility outcomes (*n* = 9), further future studies will be required to confirm these findings.

### H-Reflex responses and M-wave

As observed at rest by de Boer *et al.* (11) following 23 days of ULLS, our protocol did not modify the maximal M-wave obtained at rest, and during a weak knee extensor contraction (10% MVC). This result indicates that sarcolemmal excitability and potential action propagation were not altered by limb unloading (75,76).

In the present study, unloading increased the spinal excitability of the VM at rest, as revealed by the rise of 33% in H_MAX_/M_MAX_. Although there is no available data on knee extensors, this finding is consistent with numerous earlier studies that have shown an increase in spinal excitability of plantar flexors after 23-28 days of ULLS (7,8,12–14,29,30). Our results showed that H_MAX_/M_MAX_ obtained during a weak contraction (10% MVC) was not altered by ULLS. Since presynaptic inhibitions are already partially released during voluntary contraction, the possibility of observing an increase in the H_MAX_/M_MAX_ ratio following the ULLS is limited in this condition. However, it should be noted that the background EMG activity (i.e., the RMS of VM EMG preceding H_MAX_ stimulation, RMS/M_MAX_) was different between LS0 and LS10. To overcome that limit, the H_MAX_ was normalized by the RMS_H_ (H_MAX_/RMS_H_). The latter parameter appeared to be 2-fold greater after the 10 days of ULLS, meaning that, for a similar muscle electrical activity, the amplitude of the maximal H-reflex was increased. In light of these results, it appears that unloading induced an enhancement of H-reflex amplitude both at rest and during contraction. As shown by Delwaide (34), this enhancement could be ascribed to an increase in motoneuronal excitability and/or a decrease in presynaptic inhibition. Particularly, the latter explanation has been confirmed in an immobilization model of disuse (28). In their study, Lundbye-Jensen & Nielsen (28) showed that after immobilization of the left foot and ankle joint by a cast for 14 days, the increase in *soleus* H-reflex results from decreased presynaptic inhibition of the Ia afferents. More specifically, the authors reported a reduction in the primary afferent depolarisation mechanism (i.e., a decrease in the inhibitory activity of a spinal GABAergic interneuron) and a reduction in homosynaptic post-activation depression (i.e., a decreased quantity of neurotransmitters available in the synapse following repeated solicitation of the Ia afferents).

The present analysis showed that the decrease in muscle activation after a short period of ULLS is accompanied by an increased spinal excitability. The latter could be viewed as a compensatory adaptation, implemented by the nervous system to counterbalance the reduced capacity to activate knee extensors. However, this adaptation is not sufficient to counteract the entire cortical changes that occur during muscle disuse. Additionally, it seems that the increase in the H-reflex following ULLS is not related to the activation capacity, as reported by Seynnes *et al.* (30) on the plantar flexors after 24 days of limb unloading. Indeed, the authors did not observe a significant relationship between relative changes in amplitude of resting H-reflex and V-wave obtained during MVC over the ULLS period, indicating that the factors modulating the former had a limited effect on the latter (30). The monosynaptic T-reflex of knee extensors, evoked by patellar tendon striking, was also used as a supplementary tool to investigate the modulation of spinal circuitry following knee extensors unloading. We observed no change in the relationship between the T-reflex of the VM and the associated twitch, even if the spinal excitability was enhanced. This result suggests that other factors that affect T-reflex, including the mechanical properties of the tendon-aponeurosis and the sensitivity of the muscle spindle, have probably been altered in such a way as to mask the effect of increased spinal excitability on T-reflex (29). In contrast to our results, Seynnes *et al.* (29) reported an increase in the slope and X-intercept of the T-reflex recruitment curve. Likely, the discrepancy between the present and earlier findings may likely be ascribed to the muscle group investigated (knee extensors *vs.* plantar flexors) and the ULLS duration (10 *vs.* 23 days). However, these interpretations should be taken with caution because of the limited number of subjects (*n* = 6) that underwent the T-reflex testing procedure during the present investigation.

In the current study, neither H_LATENCY_, M_LATENCY_ nor H-M_LATENCY_ were affected by 10 days of lower limb suspension, suggesting that nerve conduction velocity through the reflex arc and synaptic delay at the spinal level were not impaired by unloading. This outcome is in contrast to those of Clark *et al.* (13) and Seynnes *et al.* (30) who found an increase in soleus M_MAX_ and H_MAX_ latencies after 28 and 24 days of ULLS respectively. More precisely, since the H-M_LATENCY_, an index of conduction velocity through the spinal reflex loop, was not changed in their studies (13,30), authors primarily attributed the slowed conduction of H-reflex and M-wave responses to the conduction velocity in the supply nerves, the branching axon terminals, and/or transmission across the neuromuscular junction (77). No change in nerve conduction through the reflex arc occurred after 10 days of ULLS for the lower limb muscles, and confirm similar evidences from Manganotti *et al*. (78) who used the bed rest model during an identical period of disuse. Further studies using ULLS protocols exceeding 10 days are needed to better understand the modulation kinetics of nerve conduction in knee extensors.

In conclusion, the present study was the first to investigate the influence of short-duration (10 days) ULLS on the neuromuscular function and spinal excitability of knee extensors. The strength loss of knee extensors, observed after 10 days of lower limb unloading, was mainly attributed to neural impairments, such as a reduction in central activation. To compensate for reduced peripheral neurosensory inputs during the ULLS protocol, the spinal excitability of the VM was enhanced at rest and during a weak knee extension (10% MVC) after the ULLS protocol. These findings are paramount since they bring out new insights into the neural adaptations that occur in the major antigravity muscle in response to muscle disuse. Understanding the physiological adaptations during ULLS can guide the development of effective interventions to improve overall health outcomes for individuals experiencing muscle inactivity. For instance, using tendon vibrations (79,80) could be relevant to stimulate Ia afferents in patients immobilised with a cast to improve spinal regulation of neural inputs (81) and maintain efficient control of force production (82,83).

## Supporting information

https://doi.org/10.6084/m9.figshare.26779975

## ACKNOWLEDGMENTS

The authors thank all volunteers who participated in the study.

## DATA AVAILABILITY

Data will be made available on reasonable request.

## AUTHOR CONTRIBUTIONS

L.L., M.V.N., A.M., G.D., F.S. and G.S. conceived and designed research; L.L., F.S and G.S performed experiments; L.L. and F.S. analyzed data; L.L., M.V.N, A.M., G.D., F.S. and G.S. interpreted results of experiments; L.L. prepared figures; L.L. drafted manuscript; L.L., M.V.N, A.M., G.D., F.S. and G.S. edited and revised manuscript; L.L., M.V.N, A.M., G.D., F.S. and G.S. approved final version of manuscript.

## SUPPLEMENTARY DATA

Supplementary data: https://doi.org/10.6084/m9.figshare.26779975

