## Supplementary material for "Effect of 10 days of unilateral lower limb suspension on knee extensors neuromuscular function and spinal excitability": https://doi.org/10.6084/m9.figshare.26779975

**TITLE:**

Université de Bourgogne - UFR STAPS

Campus Universitaire, BP 27887 DIJON

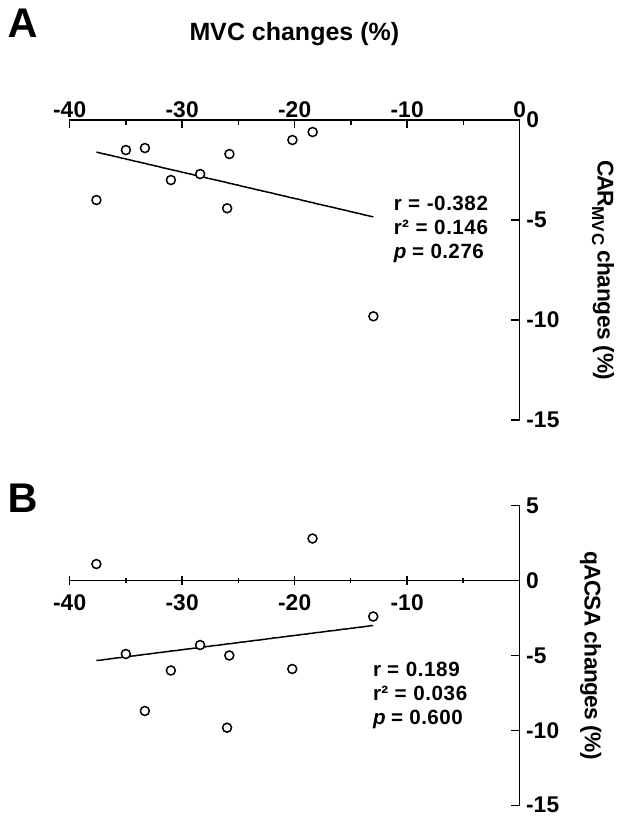

**Figure S1.** Correlations between MVC changes and changes in CAR_MVC_ (A) and qACSA (B) (*n* = 10). MVC = maximal voluntary contraction; CAR_MVC_ = central activation ratio associated with maximal voluntary contraction; qACSA = quadriceps muscle anatomical cross-sectional area.
